# A novel endosome-escaping, macrophage-targeted nanoparticle platform for miR-146a delivery with favorable in vivo biodistribution and biocompatibility

**DOI:** 10.64898/2026.02.23.707596

**Authors:** Mohammad I. Khan, Karunakaran R. Sankaran, Shaik O. Rahaman

## Abstract

Advanced nanocarrier technologies have reshaped treatment paradigms for inflammatory and degenerative disorders by facilitating cell-specific delivery of bioactive molecules, including nucleic acids. Despite this progress, therapeutic application of microRNAs (miRs) has been hindered by rapid degradation, limited stability in circulation, and suboptimal cytosolic delivery within complex biological environments. In this study, we engineered and validated a macrophage-directed lipid nanoparticle (LNP) system designed to efficiently deliver the anti-inflammatory microRNA miR-146a (MacLNP-miR146a). Mannose-functionalized LNPs were generated through a scalable lipid injection formulation approach, producing highly uniform nanoparticles with strong physicochemical integrity across diverse pH conditions and in serum-rich environments. The optimized four-lipid composition supports efficient miR-146a encapsulation, promotes endosomal escape, and enhances intracellular trafficking, leading to effective cellular uptake and favorable tissue distribution in both *in vitro* and *in vivo* models. Notably, MacLNP-miR146a demonstrates strong biocompatibility in primary cell systems and animal studies. Together, these findings position MacLNP-miR146a as a robust and translational nanotherapeutic strategy for modulating macrophage-driven inflammation, including biomaterial-associated foreign body responses and related inflammatory pathologies.

## Introduction

Engineered nanomaterials including drug delivery carriers, advanced wound dressings, and implantable biomaterials are increasingly utilized to transport therapeutic small molecules, proteins, and nucleic acids for the management of severe conditions such as cardiovascular disease, infection, chronic wounds, inflammatory bowel disorders, and autoimmune pathologies (1-4). Among nucleic acid delivery platforms, LNPs have gained particular prominence, especially following their success in clinically approved RNA-based vaccines. Conventional RNA-loaded LNPs are typically constructed from four core lipid constituents: an ionizable lipid, a structural phospholipid, cholesterol, and a polyethylene glycol (PEG)-modified lipid (5-7). These constituents perform complementary functions-ionizable lipids electrostatically associate with negatively charged RNA under acidic conditions; structural phospholipids support membrane architecture and fusion; cholesterol regulates bilayer rigidity and stability; and PEG-lipids reduce aggregation and nonspecific interactions while prolonging systemic circulation (5-10).

Despite their therapeutic promise, LNP systems are susceptible to physicochemical instability, including oxidative damage, hydrolysis, and enzymatic degradation in biological fluids, which can compromise storage stability and functional performance (11,12). To overcome these limitations and enhance miR delivery efficiency, we developed a macrophage-targeted LNP formulation encapsulating miR-146a (MacLNP-miR146a) using a defined four-lipid composition at a 40:10:48:2 molar ratio. This formulation incorporates the fusogenic helper lipid 1,2-dioleoyl-sn-glycero-3-phosphoethanolamine (DOPE) to facilitate endosomal release; the cationic lipid 1,2-dioleoyl-3-trimethylammonium-propane (DOTAP) for effective miR complexation; linoleic acid (LA) to promote membrane destabilization and intracellular delivery; and PA-PEG3-mannose to enable macrophage-selective targeting while contributing to colloidal stability (8,13).

MicroRNAs are short (18-22 nucleotide), single-stranded, noncoding RNAs that regulate gene expression post-transcriptionally by binding to complementary sequences within the 3′ untranslated regions of target mRNAs, leading to translational repression or transcript degradation (14-18). Through coordinated regulation of multiple targets, miRs function as powerful modulators of cellular phenotype, influencing activation, proliferation, differentiation, inflammatory signaling, and fibrotic remodeling (14-19). Aberrant miR expression has been implicated in diverse pathological states, including cancer, fibrosis, and the FBR to implanted biomaterials (20-24). Among inflammation-associated miRs, miR-146a is recognized as a key negative regulator of innate immune signaling (25-29). It attenuates inflammatory cascades by targeting critical mediators such as TNF receptor-associated factor 6 (TRAF6) and components of NF-κB and Toll-like receptor (TLR) pathways (30-38). Our recent work establishes miR-146a as a pivotal regulatory checkpoint in biomaterial-induced FBR (20). We observed a strong inverse correlation between miR-146a expression and inflammatory activation: increased miR-146a suppresses macrophage recruitment, foreign body giant cell (FBGC) formation, and fibrotic capsule development, whereas miR-146a deficiency exacerbates inflammatory cell accumulation and fibrotic remodeling in murine models (20). These findings highlight miR-146a as a central regulator of macrophage-driven inflammatory responses.

Therapeutically, encapsulation of miR-146a within LNPs offers multiple advantages over naked RNA delivery, including protection from nuclease-mediated degradation, enhanced intracellular trafficking, improved endosomal escape, and increased bioavailability. miR-loaded LNP systems have emerged as versatile tools capable of modulating inflammation, fibrosis, differentiation, and proliferative pathways across a range of disease contexts (38-42). Accordingly, macrophage-directed delivery of miR-146a using MacLNP-miR146a represents a promising strategy for selectively modulating macrophage-driven inflammatory responses at sites of tissue injury or biomaterial implantation.

In this work, we describe the rational design and scalable fabrication of a miR-146a-encapsulated MacLNP platform characterized by narrow particle size distribution and remarkable stability across a broad pH spectrum (2.5-8.0) and under serum-containing conditions. Using a streamlined lipid-injection and mixing approach, we generated uniform nanoparticles that support high miR encapsulation efficiency, efficient endosomal escape, and robust cellular uptake, resulting in effective biodistribution in both *in vitro* and *in vivo* models. Importantly, the MacLNP-miR146a system demonstrates strong biocompatibility in primary cells and animal studies. Together, these results support MacLNP-miR146a as a stable, scalable, and clinically adaptable nanotherapeutic platform for attenuating host inflammatory reactions and mitigating biomaterial-associated immune responses.

## Materials and methods

### Materials

To formulate LNPs and manosylated LNPs, 1,2-dioleoyl-sn-glycero-3-phosphoethanolamine (DOPE 18:1), 1,2-dioleoyl-3-trimethylammonium-propane (chloride salt) (DOTAP 18:1), 1,2-dipalmitoyl-sn-glycero-3-phospho ((ethyl-1’,2’,3’-triazole) triethyleneglycolmannose) (ammonium salt) (PA-PEG3-Mannose 16:0), 1,2-dioleoyl-sn-glycero-3-phosphoethanolamine-N-(7-nitro-2-1,3-benzoxadiazol-4-yl) (ammonium salt) (NBD PE 18:1) and 1,2-dioleoyl-sn-glycero-3-phosphoethanolamine-N-(Cyanine 5.5) were purchased from Avanti Polar Lipids, USA. D-α-Tocopherol polyethylene glycol 1000 succinate (TPGS), Linoleic acid (LA), Polyethyleneimine (PEI) and HEPES buffer were purchased from Millipore-Sigma, and all the lipids and reagents were used without further purification. Certified Low Range Ultra Agarose, Tris-Borate-EDTA (TBE) buffer, DNA Gel Loading Dye (6X), DNA ladder and Sodium Dodecyl Sulfate (SDS) were acquired from Bio-Rad. Dulbecco’s Modified Eagle Medium (DMEM), Minimum Essential Medium (MEM), Fetal Bovine Serum (FBS), Antibiotic-Antimycotic and Phosphate Buffered Saline (PBS) were purchased from Thermo Fisher Scientific. MTT (3-[4,5-dimethylthiazol-2-yl]-2,5 diphenyl tetrazolium bromide), Dimethyl sulfoxide (DMSO) and D-(+)-Mannose was purchased from Millipore-Sigma. LIVE/DEAD™ Viability Kit for mammalian cells was purchased from Invitrogen. Cell culture essentials like Dulbecco’s modified Eagle’s medium (DMEM), fetal bovine serum (FBS), antibiotic-antimycotic, and related reagents were purchased from Gibco. Ambion Pre-miR miRNA Precursor (miR-146a-5p), Pre-miR miRNA Precursor Negative Control (miR-scramble), Cy3 and Cy5.5 Dye-Labeled Pre-miR Negative Control (Cy3-miR-scramble), LysoTracker Green DND-26 and ProLong Diamond 4′,6-diamidino-2-phenylindole (DAPI) were purchased from Thermo Fisher Scientific. CD206 antibody, interleukin-4 (IL-4), granulocyte-macrophage colony-stimulating factor (GMCSF), and macrophage colony-stimulating factor (MCSF) were sourced from R&D Systems.

### MacLNP (MLNP) formulation

MacLNPs were formulated using an ethanolic injection method combined with vigorous stirring, as previously described (41). Ethanolic stock solutions of DOPE, DOTAP, PA-PEG3-Mannose were prepared in 100% ethanol, while LA was used directly in its liquid form. The lipid stocks were further diluted in 100% ethanol to achieve a final molar ratio of 40:10:48 for DOPE:DOTAP:LA respectively whereas PA-PEG3-Mannose diluted to get 2, 6 and 12 molars and coded as MacLNP2, MacLNP6 and MacLNP12. MacLNPs was prepared using above lipid combination by replacing PA-PEG3-Mannose with 2 molar TPGS. The lipids mixture (500 μl) was loaded into a Hamilton syringe and injected into 5 ml of 20 mM HEPES buffer (pH 7.4) under continuous stirring condition and maintained for 60 min. Following stirring at 500 rpm for 60 min, the solution was filtered through a 0.4 μm syringe filter to remove aggregates and obtain a homogeneous formulation. The final MacLNP suspension was stored at 4°C and used within one week.

### Fabrication of miR-146a-loaded MacLNPs and LNPs

Both MacLNP12 and LNP were freshly formulated according to the above-stated protocol and used for loading miR-146a (Ambion Pre-miR miRNA Precursor). Polyplexes were first generated using three different N/P ratios where *N* represents the moles of amine groups from PEI and *P* represents the moles of phosphate groups from miR-146a. The *N* values were set to 1, 10 and 20, while *P* was kept constant at 1. To make polyplexes, the appropriate amount of miR-146a was added to 50 μl of nuclease-free water in three different tubes marked as 1:1, 1:10, 1:20, followed by gentle mixing with a micropipette. Subsequently, the corresponding amounts of PEI, representing 1, 10, and 20 moles of *N*, were added to the respective tubes. The miR-146a/PEI mixtures were sonicated for 5 min at room temperature (RT) using a bath sonicator to form polyplexes. Simultaneously, 100 μl of MacLNP12 was taken into three different tubes and sonicated for 5 min at RT to generate empty-MacLNP12. To generate miR-146a-loaded MacLNP12, the polyplex solutions were added to the corresponding MacLNP12 suspensions and further sonicated in bath sonicator for 5 min at RT. The resulting miR-146-loaded MacLNP12 formulations were kept at RT for 1 h to allow stabilization and then ultrafiltered using 50K NMWCO ultrafiltration tube (Amicon) followed by centrifugation at 5000 rpm for 15 min to remove free or unencapsulated miR-146a. The miR-146-loaded MacLNP12 were recovered from the upper chamber of the filter tube, and the volume was adjusted to 100 μl using 20 mM HEPES buffer (pH 7.4) to obtain a final miR-146a concentration of 1000 nM in 1 mg/ml MacLNP12. Agarose gel electrophoresis was performed to confirm miR-146a loading at different N/P ratios. For this analysis, MacLNP12-miR146a samples were treated with 0.5% SDS for 5 min at a 1:1 ratio, mixed with 6x DNA loading buffer, and loaded onto a 2% agarose gel containing 0.2 μg/ml ethidium bromide. Gel was electrophoresed in 1x Tris-Borate-EDTA (TBE) buffer (Bio-Rad) at 100V for 90 min. To assess storage stability, samples were stored at 4°C for up to 8 days and analyzed by agarose gel electrophoresis. Based on loading efficiency and stability, the formulation prepared at an N/P ratio of 1:10 was selected for further characterization and downstream experiments.

### Physicochemical characterization

Empty-LNPs, miR-146a-loaded LNPs, empty-MacLNP12 and miR-146a-loaded MacLNP12 were characterized to determine their hydrodynamic size and polydispersity index (PDI) in milli-Q water using dynamic light scattering (DLS), while zeta potential was measured by phase analysis light scattering (PALS) in milli-Q water (NanoBrook Omni; Brookhaven Instruments, USA). The morphology of LNPs and miR-146a-loaded LNPs was assessed by transmission electron microscope (TEM; LVEM 5, Delong Instruments, Czech Republic).

### Stability study of miR146a encapsulated within MacLNPs and LNPs

The stability of miR-146a encapsulated in LNP, MacLNP2, MacLNP6 and MacLNP12 formulated at an N/P ratio of 1:10 was investigated at 4°C on Day 0, 7 and 14 by agarose gel electrophoresis. Moreover, stability or degradation of miR-146a within MacLNP12 was assessed at 37°C under different pH conditions using 20 mM HEPES buffer and in the presence of 1% and 10% FBS. For pH-dependent stability studies, MacLNP12-miR146a were mixed at a 1:1 ratio with 20 mM HEPES buffer adjusted to pH 2.5, 4.5, 7.4, 8.0, or 10.0 and incubated at 37°C for 1 h and 24 h. For FBS stability study, MacLNP12-miR146a was mixed with 1% and 10% mouse serum at a 1:1 ratio and incubated for 1 h at 37°C. Following incubation, all the treated samples were mixed with 0.5% SDS at a 2:1 ratio (two parts sample to one-part SDS) and kept at room temperature for 5 min to lyse the LNPs. Both SDS-treated and -untreated samples were then mixed with 6x DNA loading buffer and loaded onto a 2% agarose gel containing 0.2 µg/ml EtBr. Electrophoresis was performed for 90 min in 1x TBA buffer at 100V. Results were visualized and acquired using a gel documentation system.

### Animal maintenance and cell culture

C57BL/6 mice were purchased from The Jackson Laboratory (ME, USA). All animal experiments were conducted with protocols approved by the Institutional Animal Care and Use Committee (IACUC) at the University of Maryland College Park (Protocol No. R-OCT-24-36). Animals were housed in a pathogen-free environment under controlled temperature and humidity conditions, with food and water provided ad libitum. Primary thioglycolate-induced peritoneal-derived macrophages (MPMs) were isolated from 6-8 weeks old wild type (WT) C57BL/6 mice following our previously reported protocol (20). MPMs were cultured in DMEM supplemented with 10% FBS and 1x antibiotic-antimycotic solution. Primary mouse dermal fibroblasts (MDFs) were also isolated from the C57BL/6 mice and maintained in the minimum essential medium (MEM) media, as described in our previous reports (43-46).

### Cytotoxicity testing by MTT Assay

Cytotoxicity of MacLNPs toward MPMs was determined by MTT assay (47,48). Briefly, 2 x 10^5^ MPMs were seeded in 96-well plate in DMEM supplemented with 10% FBS and cultured for 24 h. The culture medium was then replaced with fresh complete DMEM supplemented with increasing concentrations of LNPs, MacLNP2, MacLNP6 and MacLNP12 (0, 2, 10, 15, 30, 50, and 100 μg/ml), and cells were incubated for 24 h at 37°C. Following incubation, MacLNPs/LNPs containing media were discarded from all the wells and replaced with serum free respective media containing 0.5 mg/ml MTT dye, followed by incubation for 3 h at 37°C to allow formation of purple formazan crystals by mitochondrial dehydrogenase enzymes. After incubation, the MTT-containing medium was discarded, and the formazan crystals were solubilized by adding DMSO. Absorbance was measured at 570 nm using a microplate reader, and percentage cell viability was calculated.

### LIVE/DEAD viability assay

MPMs (4 x 10^5^ cells) were seeded onto 12 mm sterile glass coverslips in 24-well plates containing their respective culture media, as mentioned above, and incubated for 48 h at 37°C (46). Following incubation, the medium was replaced with complete medium supplemented with 50 μg/ml LNP, MacNLP2, MacLNP6 and MacLNP12 and cells were further incubated for 48 h at 37°C. Untreated wells served as controls. After incubation, wells were washed twice with 1 x PBS and stained according to the manufacturer’s instruction using the LIVE/DEAD® Viability/Cytotoxicity Assay Kit (Invitrogen; L3224). Fluorescence microscopy images were obtained and processed using Fiji ImageJ software to quantify cell viability.

### Cellular uptake of NBD-labeled LNP and MacLNP

To investigate cellular uptake of LNPs and MacLNPs in MPMs, NBD-PE-tagged LNP, MacLNP2, MacLNP6 and MacLNP12 were formulated by replacing 2 mM DOPE with 2 mM NBD-PE, following the above-stated molar ratio and formulation method. The lipid mixtures were loaded into a Hamilton syringe and injected into 20 mM HEPES buffer under continuous stirring, followed by incubation in the dark for 1 h. MPMs (4 x 10^5^ cells) were seeded onto 12 mm sterile glass coverslips in 24-well plates containing their respective culture media and incubated for 48 h at 37°C. Following incubation, the medium was discarded, and fresh complete medium containing 50 μg/ml NBD-PE-tagged MacLNP2, MacLNP6, MacLNP12 and LNP was added, and cells were further incubated for 1 h at 37°C. Following treatment, cells were washed twice with ice-cold 1 x PBS to halt further internalization. Cellular uptake was visualized using a fluorescence microscope. Acquired images were processed using Fiji ImageJ software.

### Mannose competitive uptake

MPMs (4 x 10^5^ cells) were seeded onto 12 mm sterile glass coverslips in 24-well plates in DMEM with 10% FBS and incubated for 48 h at 37°C. Following incubation, the medium was discarded, and fresh complete medium containing 50 mM D-(+)- mannose was added and incubated for 1h at 37°C. After 1h, mannose containing media was discarded and MPMs were washed twice with 1x PBS. Then 50 μg/ml NBD-PE-tagged MacLNP12 and LNP was added, and cells were further incubated for 1 h at 37°C. Following treatment, cells were washed twice with 1 x PBS. MacLNP12 and LNP cellular uptake in the presence and absence of mannose was visualized using a fluorescence microscope. Acquired images were processed using Fiji ImageJ software.

### Immunoblotting

Presence of mannose receptor on tested cells were confirmed by western blotting. BMDM, MPM, MDF and VIC were grown in their respective media in 60 mm tissue culture dish. At 90% confluency, cells were harvested in RIPA (Radioimmunoprecipitation Assay) buffer containing both protease and phosphatase inhibitors and whole cell lysate (WCL) prepared. Protein concentration on WCL was estimated using BCA assay kit (Thermo fisher). Presence of CD206 was investigated by western blotting in all the above-mentioned cells and compared with anti-Actin IgG (Cell Signaling Technology; 4970).

### Uptake of Cy3-labeled scrambled-miR-146a-loaded MacLNPs and LNPs

To assess the transfection efficiency and intracellular stability of miR-146a over time, Cy3-labeled scrambled-miR-146a was loaded into MacLNP12 and LNP at an N/P ratio of 10 using the protocol described above and used immediately. MPMs (4 x 10^5^ cells) were seeded onto 12 mm sterile glass coverslips in 24-well plates containing their respective culture media and incubated for 48 h at 37°C. Following incubation, the culture medium was replaced with fresh complete medium, and cells were transfected with MacLNP12-Cy3-Scr-miR146a and LNP-Cy3-Scr-miR146a at a final concentration of 50 nM. Cells were incubated for 1, 2 and 24 h at 37°C. After incubation, the medium was removed, and cells were washed twice with 1 x PBS and fixed with 4% paraformaldehyde, followed by additional washes with 1 x PBS. Coverslips were mounted using a DAPI-containing solution and allowed to cure overnight at room temperature in the dark. The presence of Cy3-Scr-miR146a loaded MacLNP12 and LNP and nuclei was visualized using a fluorescence microscope. Images were captured and processed using Fiji ImageJ software.

### Endo/lysosomal escape study

MPMs (4 x 10^5^ cells) were seeded onto 12 mm glass coverslips in 24-well plates and incubated at 37°C for 48 h. The culture medium was then replaced with fresh complete medium containing 50 nM LysoTracker™ Green DND-26 (Invitrogen) to stain lysosomes. Cells were subsequently treated with 50 nM MacLNP12-Cy3-Scr-miR146a and incubated for 2 h at 37°C. For the 24 h time-point study, cells were first treated with 50 nM MacLNP12-Cy3-Scr-miR146a, and at 22 h post-treatment, 50 nM LysoTracker was added and incubated for an additional 2 h at 37°C. After incubation, the medium was discarded, and cells were washed twice with 1 x PBS, fixed with 4 % paraformaldehyde, and washed again with 1 x PBS. Glass coverslips were carefully removed and inverted on rectangular glass slides containing a DAPI-containing mounting medium, then kept in the dark overnight to allow solidification. Images were acquired using a confocal microscope (FLUOVIEW FV3000) with DAPI (blue), FITC (green), and TRITC (red) filter channels to visualize nuclei, lysosomes, and Cy3-miR-Scr-loaded MacNLP12, respectively. Image analysis was performed using Fiji ImageJ software.

### In vivo cytotoxicity analysis

To investigate in vivo biocompatibility, 6-8 weeks old WT C57BL/6 mice were selected and divided in three groups (n = 3 per group): HEPES, MacLNP12 (MLNP12), and MacLNP12-miR146a. MacLNP12 was injected intravenously via the tail vein at a dose of 1 mg/kg body weight. In MacLNP12-miR146a group, 50 uL of MacLNP12 containing 0.05 nmol of miR-146a were injected. Control mice received 50 μl of HEPES buffer only. All the mice were housed in a sterile environment for 7 days. After the incubation period, mice were euthanized, and blood was collected for further analysis. Serum was used to assess hepatic function markers - aspartate aminotransferase (AST) and alanine transaminase (ALT); kidney function markers - blood urea nitrogen (BUN) and creatinine; and basic metabolic electrolytes - calcium, bicarbonate, chloride, and potassium. Whole blood was used to assess hematological parameters including hemoglobin (HGB), hematocrit (HCT), white blood cells (WBC) count, neutrophils, lymphocytes, Monocytes, platelet count and estimate. Collected samples were sent to IDEXX BioAnalytics (North Grafton, MA, USA) for analysis. The resulting data were analyzed and plotted using GraphPad Prism.

### Analysis of subcutaneous retention and systemic biodistribution of MacLNP12-Cy3-Scr-miR146a *in vivo*

Skin tissue uptake, distribution, and stability of Cy3-tagged scrambled-miR-146a-loaded in MacLNP12 was evaluated in 6-8-weeks-old WT mice using IVIS. Dorsal skin was shaved to minimize autofluorescence interference and 50 uL MacLNP12-Cy3-Scr-miR146a (0.05 nmol) was injected subcutaneously. IVIS images were collected at 0 h, and 3 h post-injection, and subsequently on days 1, 4, and 7. An additional cohort of mice (n = 3 per group) was euthanized on day 4 post-injection, and organs (skin, liver, lungs, kidneys, spleen and heart) were harvested, placed in sterile petri dishes, and imaged using an IVIS spectrum. All images were analyzed using LivingImage software.

### Analysis of *in vivo* delivery and uptake of MacLNP12-Cy5.5-Scr-miR146a in major organs in mice

Eight-week-old ApoE^-/-^ (C57BL/6 background) mice were purchased from The Jackson Laboratory and housed as per IACUC guideline in the UMD animal facility. Mice were maintained on a high-fat diet for 6 weeks, after which the uptake and biodistribution of Cy5.5-tagged MacLNP12-Scr-miR146a were evaluated. MacLNP12-Cy5.5-Scr-miR146a were formulated following the above-mentioned MacLNP12 formulation method. The molar ratio of lipid components (DOPE:DOTAP:LA:PA-PEG3-Mannose) was 40:10:48:12, respectively. Cy5.5-PE (1 μg/µl) was mixed with the lipid mixture, loaded into a Hamilton syringe, and injected into 20 mM HEPES buffer under stirring, followed by incubation in the dark for 1 h. The final Cy5.5 concentration was adjusted to 10 μg/ml, which provided optimal excitation and emission as per manufacturer’s specifications. MacLNP12-Cy5.5-Scr-miR146a (50 μl containing 10 μg/ml Cy5.5) were intravenously injected (n = 3 per group), and mice were euthanized 3 h post-injection, and organs were harvested for analysis. The heart with the aortic arch and the full abdominal aorta were carefully collected and placed in sterile petri dishes. Additional major organs, including lungs, liver, kidneys, and spleen, were also collected and placed in separate petri dishes. Uptake and biodistribution of MacLNP12-Cy5.5-Scr-miR146a in all harvested organs were visualized using an IVIS and analyzed by LivingImage software.

### Quantitative real-time RT-PCR (RT-qPCR)

Bone marrow-derived macrophages (BMDMs) were prepared and cultured according to our previously published protocol (20). Approximately 2 x 10^6^ BMDMs were seeded in 60-mm culture dishes in DMEM supplemented with 10% FBS and incubated for 24 h at 37°C. After cell adhesion, the medium was replaced with fresh complete DMEM. miR-146a and scrambled-miR-146a were loaded into MacLNP12 and LNP at an N/P ratio of 10 prior to use. BMDMs were treated with 50 nM MacLNP12-miR146a, MacLNP12-Scr-miR146a, LNP-miR146a and LNP-Scr-miR146a in complete DMEM and incubated for 24 h at 37°C. Following treatment, the medium was discarded, cells were washed twice with 1 x PBS, and cells were harvested form each dish. Total RNAs and microRNAs were extracted from BMDMs according to the manufacturer’s instructions (Qiagen), followed by treatment with RNase-free DNase. The concentration and purity of the extracted RNAs and microRNAs were determined using a Nanodrop-2000 system (Thermo Fisher Scientific).

To evaluate miR-146a expression, cDNA was synthesized using the miRCURY LNA SYBR Green PCR Kit according to the manufacturer’s instructions (Qiagen) using a SimpliAmp Thermal Cycler (Applied Biosystem). RT-qPCR was then performed using the miRCURY LNA SYBR Green PCR Kit with primers for hsa-miR-146a-5p (YP00204688) and SNORD68 (YP00203911), following the manufacturer’s instructions (Qiagen). Amplification was carried out for 40 cycles using a CFX96 cycler (Bio-Rad). Threshold cycle (Ct) values for hsa-miR-146a-5p were normalized to the average Ct value of SNORD68, and relative miR-146a expression was calculated using the ΔΔCt method. In addition, expression of TRAF6 was assessed in MacLNP12-miR146a, LNP-Scr-miR146a, and LNP-miR146a-treated BMDMs. cDNA synthesized using iScript Advance cDNA Kit (172-5037) and RT-qPCR was performed using the SsoAdvanced Universal SYBR Green Supermix (172-5270) and primers for TRAF6 (qMmuCEP0054569) and GAPDH (qMmuCID0018612), according to the manufacturer’s instructions (Bio-Rad). Reactions were run for 40 cycles on a CFX96 cycler (Bio-Rad). Ct values of target genes were normalized to GAPDH, and relative gene expression was calculated using the ΔΔCt method.

### Declaration of generative AI and AI-assisted technologies in the writing process

During the preparation of this work the author(s) used ChatGPT 5.2 in order to improve readability and perform proofreading. After using this tool/service, the author(s) reviewed and edited the content as needed and take(s) full responsibility for the content of the publication.

## Results

### Formulation and physicochemical characterization of nanomaterials

MacLNPs and LNPs were formulated using an ethanol injection method, followed by miR-146a loading via a controlled sonication method. The resulting formulations were characterized to evaluate for hydrodynamic diameter, PDI, and surface charge (zeta potential). Obtained results are presented in Figure 1A. DLS analysis revealed that empty LNPs and empty MacLNP12 particles exhibited mean diameters of 163.79 ± 7.69 nm and 148.78 ± 9.16 nm, respectively. Following miR-146a encapsulation, particle sizes measured 182.93 ± 3.38 nm for LNP-miR146a and 151.56 ± 6.67 nm for MacLNP-miR146a (Figure 1B). Importantly, miR loading did not substantially alter the PDI values for either formulation, indicating that RNA incorporation did not significantly impact particle dispersity or homogeneity. Moreover, Surface charge analysis demonstrated a marked shift in zeta potential after miR encapsulation. The initially negative surface charge of empty LNPs (−21.42 ± 4.51 mV) and empty MacLNP12 (−30.59 ± 3.46 mV) became less negative upon miR-146a loading, measuring -6.21 ± 1.25 mV and -4.84 ± 1.19 mV, respectively (Figure 1C). This reduction in negative charge is consistent with electrostatic complexation between the lipid components and the miR cargo. Loading efficacy of miR-146a in LNPs and MacLNP were observed to be ∼ 95 and 92% respectively (Figure 1A). TEM was used to examine nanoparticle morphology. Representative images (Figure 1D-E) confirmed that both empty and miR-loaded MacLNP12 particles maintained a spherical architecture within the nanoscale size range. No discernible morphological alterations were observed following miR-146a incorporation. Higher-magnification images further revealed the presence of internal polyplex structures and demonstrated particle uniformity consistent with the size distribution profiles obtained from DLS measurements.

**Figure 1.**
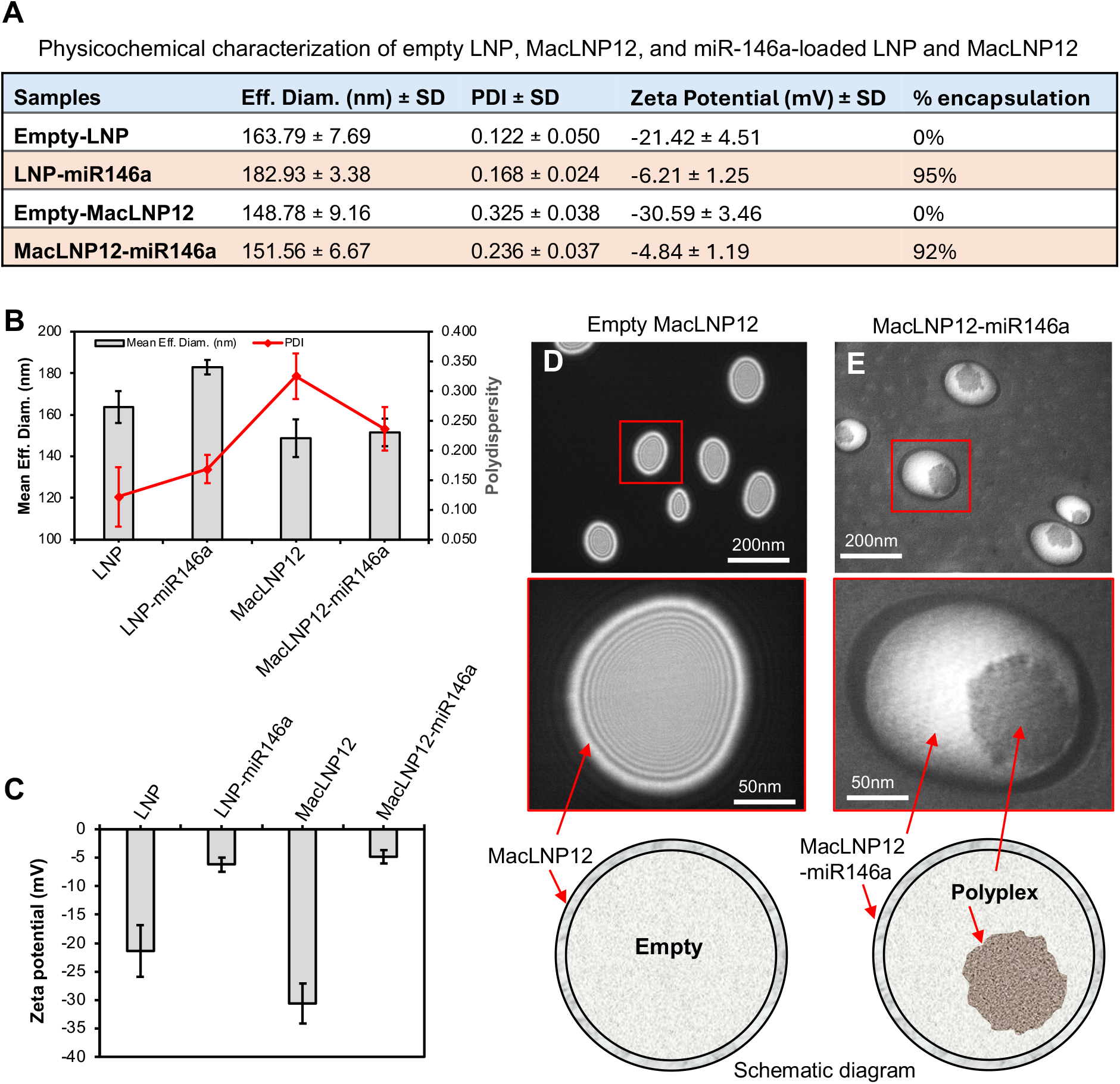
Physicochemical characterization of nanomaterials. **(A)** The table summarizes the physicochemical properties of empty-LNP, empty-MacLNP12, LNP-miR146a, and MacLNP12-miR146a, including effective diameter, PDI, zeta potential, and encapsulation efficiency. The characteristic properties of the LNPs were determined using multiple analytical techniques, including **(B)** DLS, and **(C)** PALS. **(D and E)** The morphology of empty-MacLNP12 and MacLNP12-miR146a was evaluated by TEM. All formulations were prepared in triplicate, and data are presented as the mean ± standard error of the mean.

Agarose gel electrophoresis was performed to evaluate miR-146a encapsulation within MacLNP12 prepared at three different nitrogen-to-phosphate (N/P) ratios (1:1, 1:10, and 1:20). Analysis of ultrafiltrates revealed detectable free miR-146a at the 1:1 ratio, whereas no unbound miR was observed at N/P ratios of 1:10 or 1:20, indicating more complete complexation at higher ratios (Figure 2A). MacLNP12-miR146a formulations generated at these N/P ratios were subsequently treated with 0.5% SDS on Day 0 to disrupt particles and release encapsulated RNA prior to gel electrophoresis. Clear miR-146a bands were observed for the 1:1 and 1:10 formulations, while minimal signal was detected for the 1:20 ratio (Figure 2B). To assess short-term stability, the same samples were stored at 4°C for 7 days, followed by SDS treatment and gel analysis. No appreciable loss or degradation of miR-146a was detected in the 1:1 and 1:10 groups compared to Day 0 (Figure 2C). Based on overall encapsulation and release profiles, the 1:10 N/P ratio was selected as optimal.

**Figure 2.**
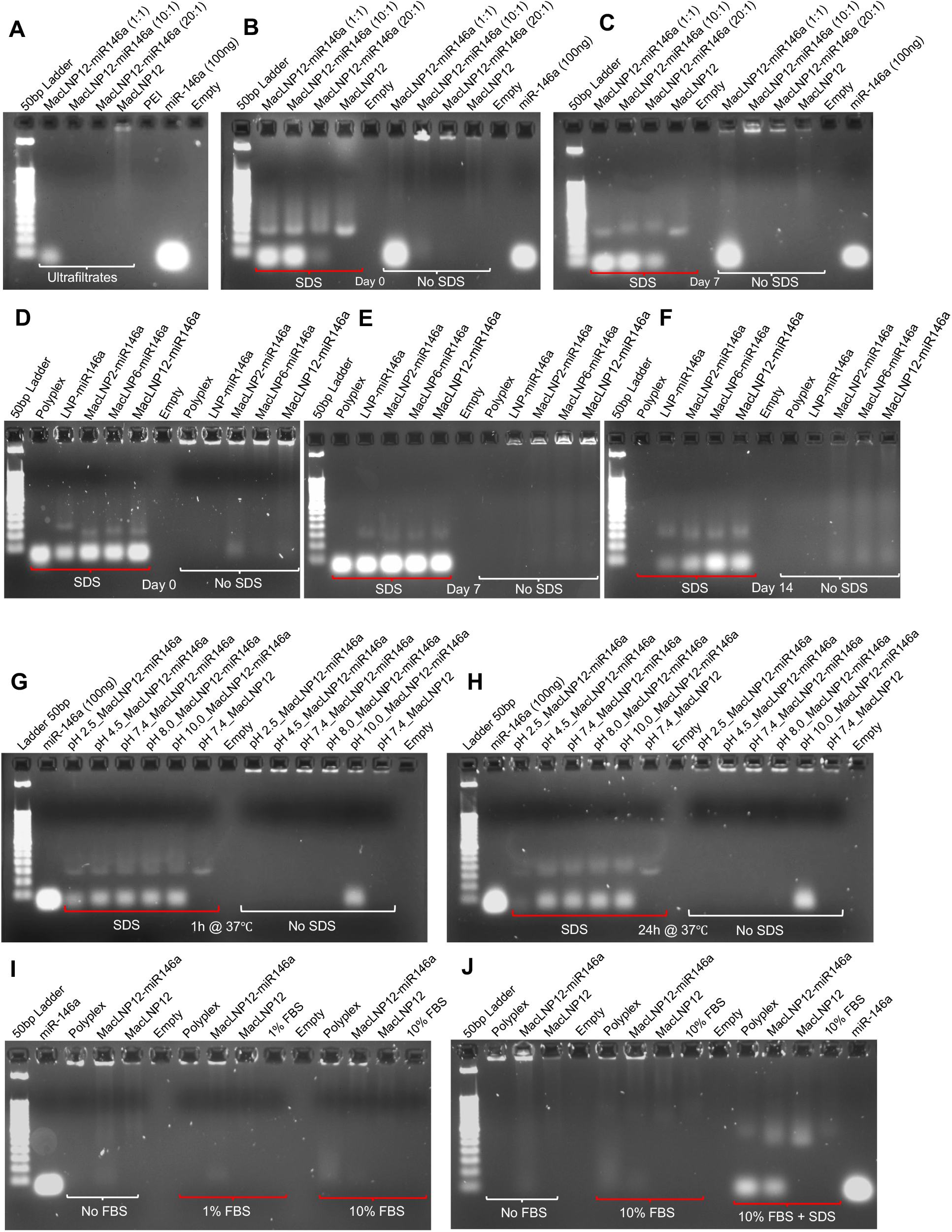
Loading of miR-146a, and stability and encapsulation efficiency in MacLNPs. **(A)** MacLNP12-miR146a was formulated using three different N/P ratios, and ultrafiltrates were analyzed by agarose gel electrophoresis to assess unencapsulated miR-146a. **(B)** MacLNP12-miR146a was treated with SDS on day 0 and subjected to agarose gel electrophoresis to determine the optimum N/P ratio. **(C)** The stability of miR-146a at 4°C was evaluated over 8 days, and SDS-treated samples were analyzed by agarose gel electrophoresis. (**D-F**) Stability of miR-146a encapsulated in LNP, MacLNP2, MacLNP6 and MacLN12 was assessed by agarose gel electrophoresis after 0, 7, and 14 days of incubation at 4°C. **(G and H)** Stability of MacLNP-loaded miR-146a under different pH conditions was checked after 1 h and 24 h of incubation. **(I and J)** Stability of miR-146a in MacLNP12 was evaluated in the presence of 1% and 10% FBS at 37°C and analyzed by agarose gel electrophoresis after 1 h of incubation.

Using this ratio, miR-146a encapsulation and stability were further compared across LNP, MacLNP2, MacLNP6, and MacLNP12 formulations. Day 0 gel analysis confirmed successful miR incorporation in all nanoparticle types (Figure 2D). After 7 days at 4°C, intact miR-146a bands remained visible in all formulations (Figure 2E). However, following 14 days of storage, diminished band intensity was observed for LNP and MacLNP2, whereas MacLNP6 and MacLNP12 retained strong and distinct miR-146a bands (Figure 2F). These findings indicate that while all formulations protect miR-146a from rapid degradation for at least 7 days, MacLNP6 and MacLNP12 provide superior stability during extended storage.

The physicochemical robustness of MacLNP12-miR146a was further examined under stress conditions. Incubation at 37°C across a range of pH values demonstrated particle destabilization at pH 10, as evidenced by the appearance of miR-146a bands on gels without SDS treatment after both 1 and 24 hours (Figure 2G, 2H). Following 1 hour of incubation, SDS-treated samples showed detectable miR-146a at all pH values, although band intensity was reduced at pH 2.5. After 24 hours, miR-146a bands were no longer detectable at pH 2.5, suggesting nanoparticle disruption under highly acidic conditions.

Serum stability was also evaluated by incubating MacLNP12-miR146a with 1% and 10% FBS at 37°C for 1 hour. Agarose gel analysis showed miR-146a bands only after SDS treatment, whereas untreated samples displayed no detectable free RNA, confirming effective protection against serum-associated degradation (Figure 2I, 2J).

Collectively, these data demonstrate efficient miR-146a encapsulation, sustained protection under storage and physiological conditions, and enhanced stability in MacLNP6 and MacLNP12 formulations. The results validate the robustness of the lipid injection and sonication-based preparation method for generating stable and homogeneous macrophage-targeted miR-146a-loaded nanoparticles.

### Excellent biocompatibility of MacLNPs in primary macrophages

Evaluation of cytocompatibility is essential to determine the translational suitability of macrophage-targeted MacLNPs. To assess cellular tolerance, primary mouse peritoneal macrophages (MPMs) were exposed to MacLNP2, MacLNP6, and MacLNP12 across increasing concentrations for 24 hours. All formulations demonstrated favorable biocompatibility profiles. At a concentration of 50 μg/ml, cell viability remained approximately 80% for MacLNP2 and around 75% for both MacLNP6 and MacLNP12 (Figure 3A). Maintaining greater than 75% viability in primary mammalian cells supports the suitability of these formulations for biomedical applications. To further validate these findings, live/dead staining was performed after 48 hours of exposure to 50 μg/ml of LNP, MacLNP2, MacLNP6, or MacLNP12. Fluorescence imaging revealed predominantly viable (green) cells with only a small fraction of nonviable (red) cells across all treated groups, comparable to untreated controls (Figure 3B). Together, these results demonstrate that both non-targeted LNPs and macrophage-targeted MacLNP formulations exhibit strong cytocompatibility in primary macrophages. Based on these findings, 50 μg/ml was selected as the working concentration for subsequent *in vitro* experiments.

**Figure 3.**
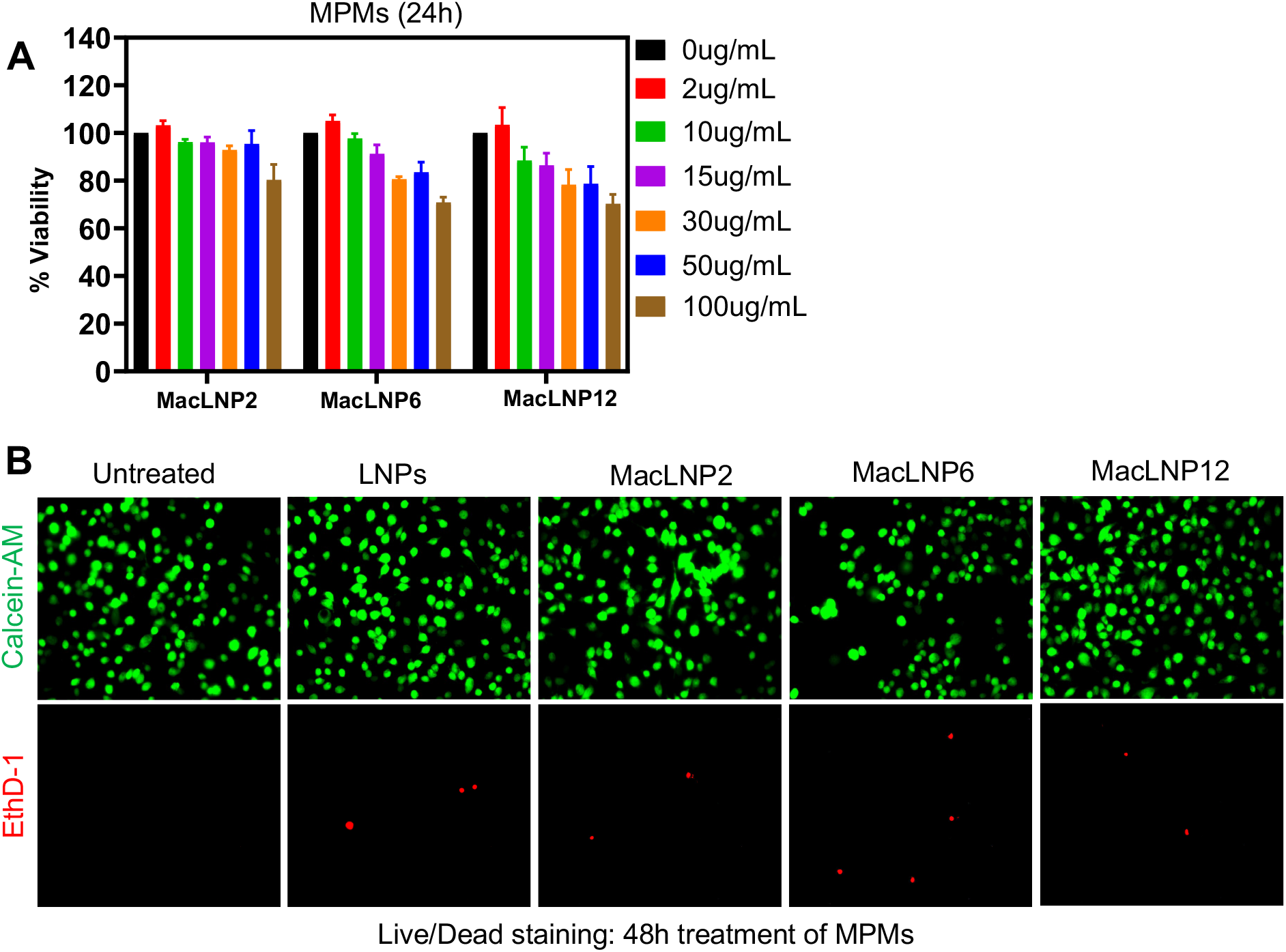
In vitro biocompatibility assessment of MacLNPs in primary macrophages. **(A)** Cytotoxicity of MacLNPs was examined using an MTT assay in MPMs following 24 h incubation with increasing concentrations of MacLNP2, MacLNP6 and MacLNP12. **(B)** Cell viability of MPMs was further checked using a LIVE/DEAD assay by fluorescence microscope after 48 h treatment with 50 µg/ml of LNP, MacLNP2, MacLNP6 and MacLNP12. All experiments were performed in triplicate, and data are presented as mean ± standard error of the mean.

### Efficient intracellular uptake of MacLNPs into primary macrophages

Fluorescence microscopy was used to examine internalization of NBD-PE-labeled LNP, MacLNP2, MacLNP6, and MacLNP12 in primary macrophages (MPMs) (Fig. 4A). Uptake of NBD-tagged MacLNP2, MacLNP6, and MacLNP12 progressively increased and was consistently greater than that observed with non-targeted NBD-LNP. Notably, cellular fluorescence intensity rose in parallel with increasing PA-PEG3-mannose incorporation, indicating effective engineering of mannose-directed, macrophage-targeted nanoparticles. To validate receptor-mediated uptake, MPMs were treated with NBD-MacLNP12 or NBD-LNP in the presence or absence of free mannose (Fig. 4B-C). In the absence of mannose, both formulations were internalized; however, MacLNP12 produced substantially higher fluorescence. Mannose pretreatment markedly reduced MacLNP12 binding and uptake, supporting a CD206-dependent mechanism. Consistently, expression of the mannose receptor (CD206) in macrophages was confirmed by immunoblotting (Fig. 4D).

**Figure 4.**
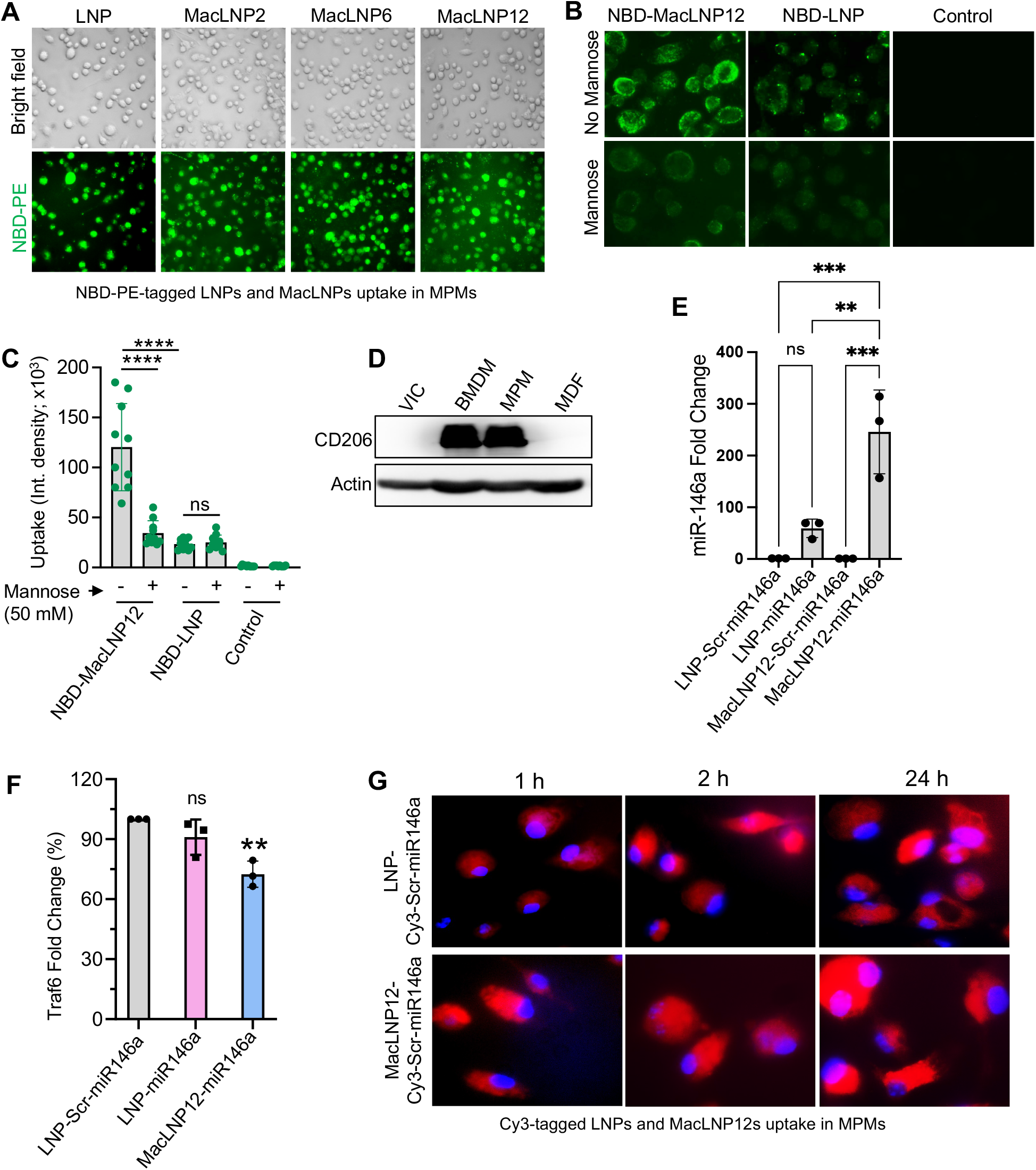
Internalization of MacLNPs by primary macrophages, receptor-mediated macrophage targeting, and efficient intracellular delivery of miR-146a. **(A)** Uptake of LNPs and MacLNPs in MPMs was assessed using NBD-PE labeling. **(B)** Inhibition of uptake in MPMs was evaluated in the presence of mannose, and **(C)** fluorescence intensity was quantified. **(D)** Immunoblotting was done to confirm CD206 expression in different cell types. (**E**) Fold-change in miR-146a expression in BMDM was determined by RT-qPCR analysis. (**F**) Expression level of Traf6 in BMDMs were determined by RT-qPCR analysis. **(G)** Uptake of LNP-Cy3-Scr-miR146a and MacLNP12-Cy3-Scr-miR146a was evaluated in MPMs. All experiments were performed in triplicates. Representative images are shown, and quantitative data are presented as mean ± standard error of the mean (SEM). Statistical significance was determined using one-way ANOVA and is denoted as **p < 0.01, ***p < 0.001, and ****p<0.0001; NS, not significant.

miR-146a expression was quantified in BMDMs 24 h after exposure to LNP-Scr-miR146a, LNP-miR146a, MacLNP-Scr-miR146a, or MacLNP12-miR146a. MacLNP12-miR146a induced an approximately fivefold greater increase in miR-146a levels compared with LNP-miR146a (Fig. 4E). Functional activity of the delivered miRNA was assessed by measuring TRAF6, a validated miR-146a target. Traf6 expression decreased by ∼30% in MacLNP12-miR146a–treated cells relative to scrambled controls (Fig. 4F), confirming successful intracellular delivery and target suppression.

Time-dependent uptake kinetics were further evaluated using Cy3-labeled scrambled miR-146a encapsulated in MacLNP12. In MPMs, intracellular fluorescence was evident as early as 1 h, with near-maximal uptake achieved rapidly and maintained through 24 h (Fig. 4G). No significant loss of Cy3 signal at 24 h (37°C) was observed, suggesting intracellular stability of the formulation. In contrast, uptake of LNP-Cy3-Scr-miR146a occurred more gradually, reaching complete internalization only after 24 h. Collectively, these findings demonstrate efficient and selective macrophage uptake of miR-146a-loaded MacLNPs, supporting their utility as targeted carriers for intracellular miR delivery.

### Endo/lysosomal escape of MacLNP12-miR146a in primary macrophages

Effective escape from endosomal and lysosomal compartments is critical for LNP-based nucleic acid delivery, as it protects cargo from degradation and enables release into the cytosol. To assess intracellular trafficking and escape efficiency, MacLNP12-Cy3-miR146a was administered to MPMs, and its localization was examined by confocal microscopy (Fig. 5A-B). At 2 h post-treatment, strong colocalization of MacLNP-Cy3-Scr-miR146a (red) with lysosomal markers (green) was detected, indicating initial endo/lysosomal sequestration. By 24 h, this overlap was substantially diminished, with clear spatial separation between red and green fluorescence signals (Fig. 5A). Quantitative colocalization analysis (yellow signal) demonstrated an approximately 70% decrease at 24 h relative to 2 h (Fig. 5B), consistent with progressive lysosomal escape. Together, these findings indicate that macrophage-targeted MacLNP-miR146a efficiently exits endo/lysosomal compartments following internalization, supporting its effectiveness as a cytosolic delivery vehicle for therapeutic nucleic acids.

**Figure 5.**
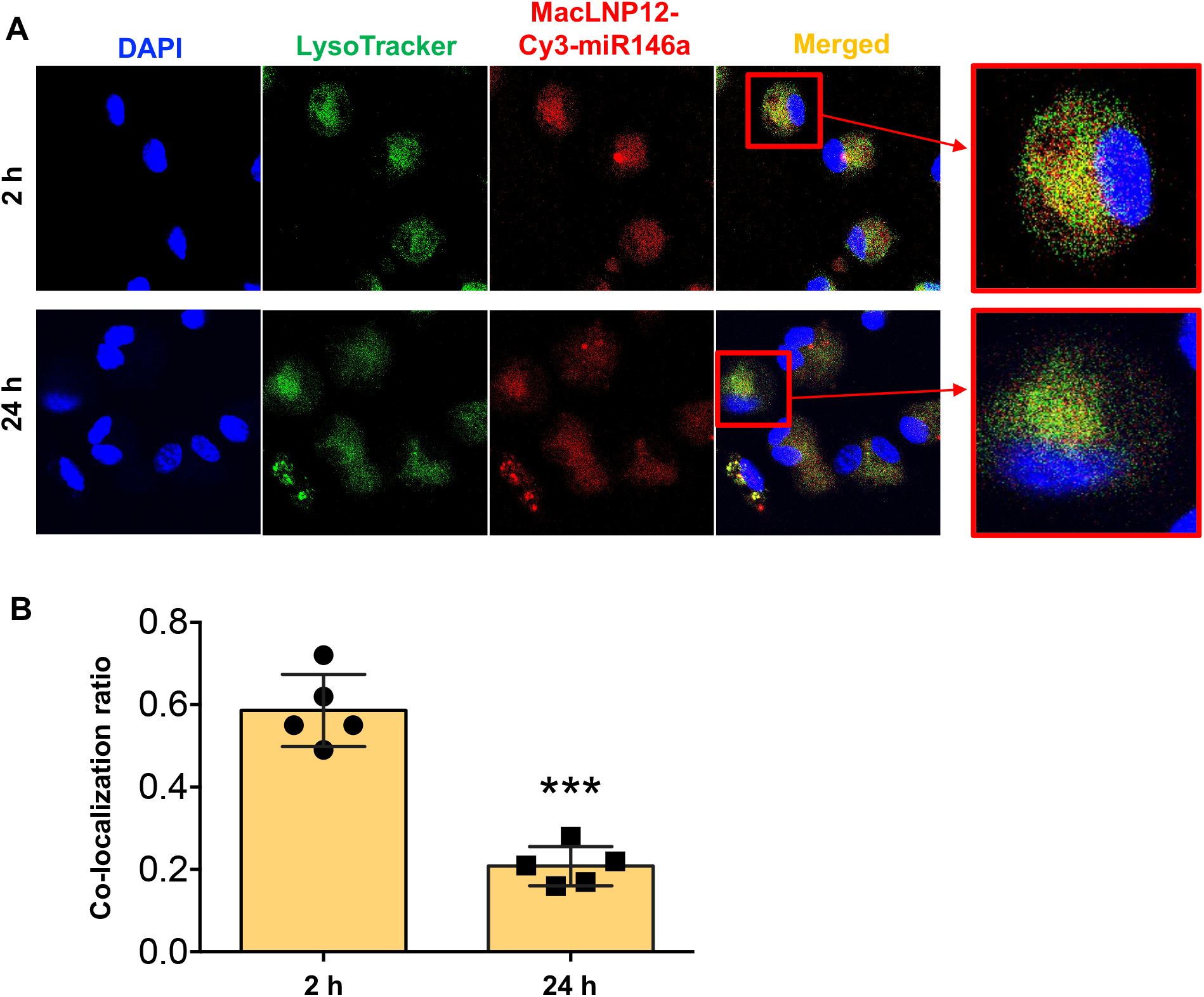
Efficient endo/lysosomal escape of MacLNP12-miR-146a. **(A)** Endo/lysosomal escape of MacLNP-Cy3-Scr-miR146a was evaluated in MPMs after 2 h and 24 h of treatment. **(B)** Escape efficiency was quantified by calculating the colocalization ratio based on the overlap of red (Cy3-Scr-miR146a) and green (lysosomal) fluorescence signals, which appear yellow in merged images. All experiments were performed in triplicate. Representative images are shown, and quantitative data are presented as mean ± standard error of the mean. Statistical significance was determined using a t-test and is denoted as ***p < 0.001.

### MacLNP-miR146a demonstrates extended in vivo stability, limited off-target biodistribution, and durable retention at the injection site

The therapeutic efficacy of MacLNP12-miR146a depends on its in vivo persistence and spatial distribution, particularly for applications requiring sustained local delivery. To characterize these parameters, we evaluated nanoparticle retention and biodistribution in a subcutaneous dorsal skin injection model in wild-type mice. Following administration of MacLNP12-Cy3-Scr-miR146a, fluorescence was monitored longitudinally by IVIS imaging. Strong epifluorescent signals were observed immediately after injection (0 h), at 3 h, and on day 1, localized within a defined radius at the injection site (Fig. 6A). Although signal intensity progressively declined, fluorescence remained detectable through day 4. Quantification of total radiant efficiency demonstrated a time-dependent decrease in signal, with a marked reduction from day 0 to day 7 (Fig. 6B).

**Figure 6.**
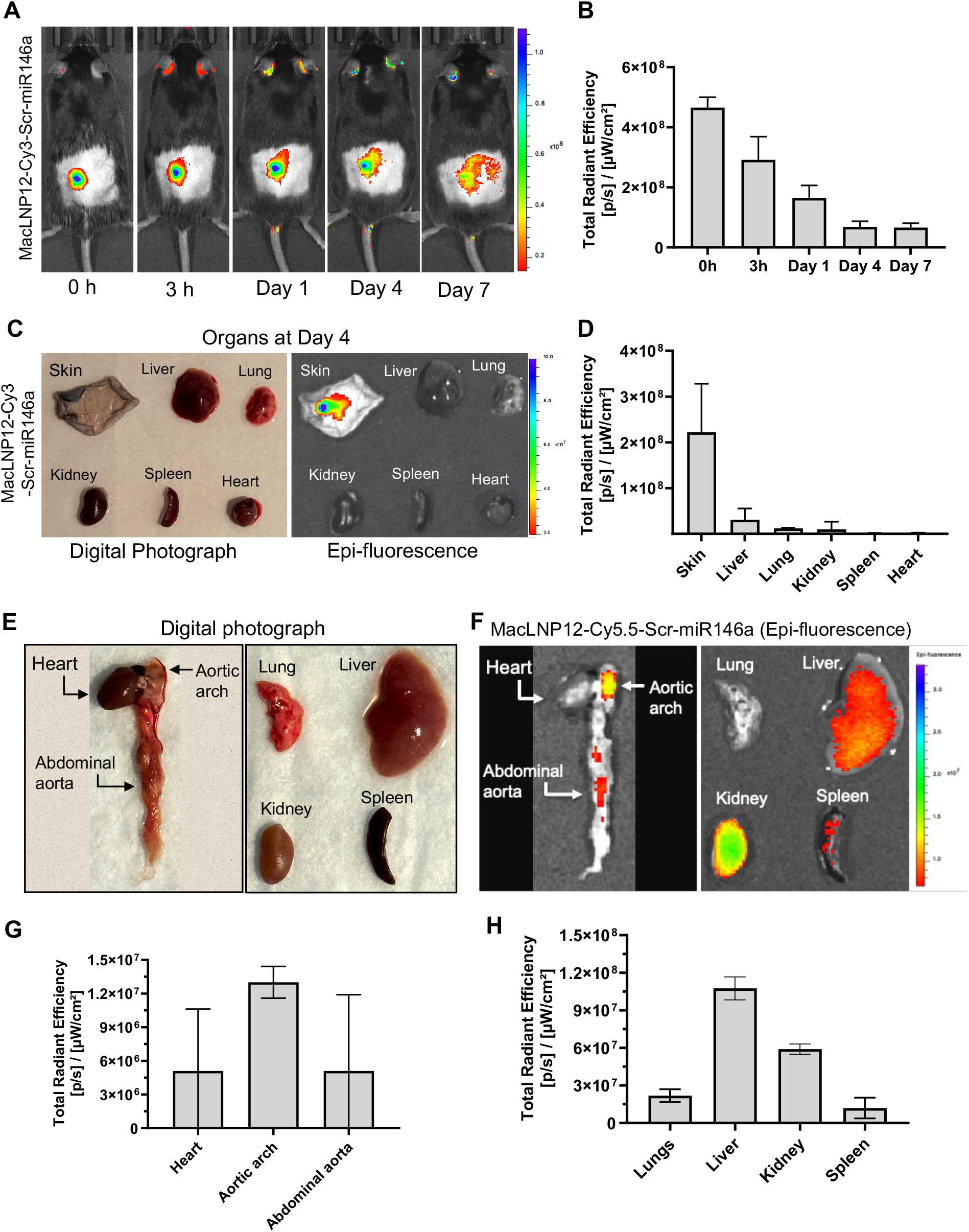
In vivo stability and tissue distribution of miR-146a-loaded MacLNP12. **(A)** In vivo uptake and distribution of MacLNP-Cy3-Scr-miR146a in skin tissue were monitored by IVIS imaging at 0 h, 3 h, day 1, day 4, and day 7 post-injection. **(B)** Quantification of total radiant efficiency of MacLNP12-Cy3-Scr-miR146a from IVIS images. **(C)** On day 4, potential leaching and migration of MacLNP-Cy3-Scr-miR146a to major organs, including skin, liver, lungs, kidneys, spleen, and heart, were evaluated. **(D)** Quantification of total radiant efficiency in each organ was quantified based on IVIS analysis. Systemic biodistribution of Cy5.5-labeled MacLNP12 was further assessed following intravenous administration in hyperlipidemic ApoE□/□ mice. Representative digital images and epifluorescence of major organs are shown **(E, F)**, and fluorescence intensity was quantified by IVIS to determine total radiant efficiency **(G, H)**. All experiments were performed in triplicates, and data are presented as the mean ± standard error of the mean.

Ex vivo imaging performed on day 4 further assessed tissue distribution in skin and major organs, including liver, lung, kidney, spleen, and heart (Fig. 6C). Fluorescence was largely restricted to the injection site, with no appreciable evidence of systemic redistribution. Radiant efficiency measurements confirmed significantly greater signal intensity in the skin relative to other organs, indicating strong local retention of MacLNP12-Cy3-Scr-miR146a (Fig. 6D).

To evaluate systemic biodistribution, MacLNP12-Cy5.5-Scr-miR146a was administered intravenously to hyperlipidemic ApoE□/□ mice. Ex vivo imaging of excised organs revealed preferential accumulation within the aortic arch and abdominal aorta, whereas minimal fluorescence was detected in the heart (Fig. 6E-F). Quantitative analysis showed significantly higher radiant efficiency in the abdominal aorta compared to the aortic arch and heart (Fig. 6G). As anticipated for nanoparticle clearance, the liver exhibited the strongest overall fluorescence signal, consistent with hepatic uptake (Fig. 6H).

Taken together, these findings demonstrate that targeted MacLNP-miR146a displays durable in vivo stability, limited off-target organ distribution, and sustained localization at the administration site, supporting its utility for both localized and vascular-targeted therapeutic delivery.

### MacLNP12-miR146a exhibits no detectable toxicity in vivo

The in vivo cytotoxicity of MacLNP12 and MacLNP12-miR146a was evaluated to assess systemic biocompatibility. Hepatic function was examined by measuring serum AST and ALT levels. AST levels were approximately 300 U/L in both HEPES- and MacLNP12-treated mice, whereas a marked reduction was observed in the MacLNP12-miR146a group, with levels near 100 U/L (Figure 7A). ALT levels ranged from 30 to 70 U/L across all groups, remaining within the normal physiological range for healthy wild-type mice (Figure 7B). These findings indicate preserved hepatic function following MacLNP12 administration.

**Figure 7.**
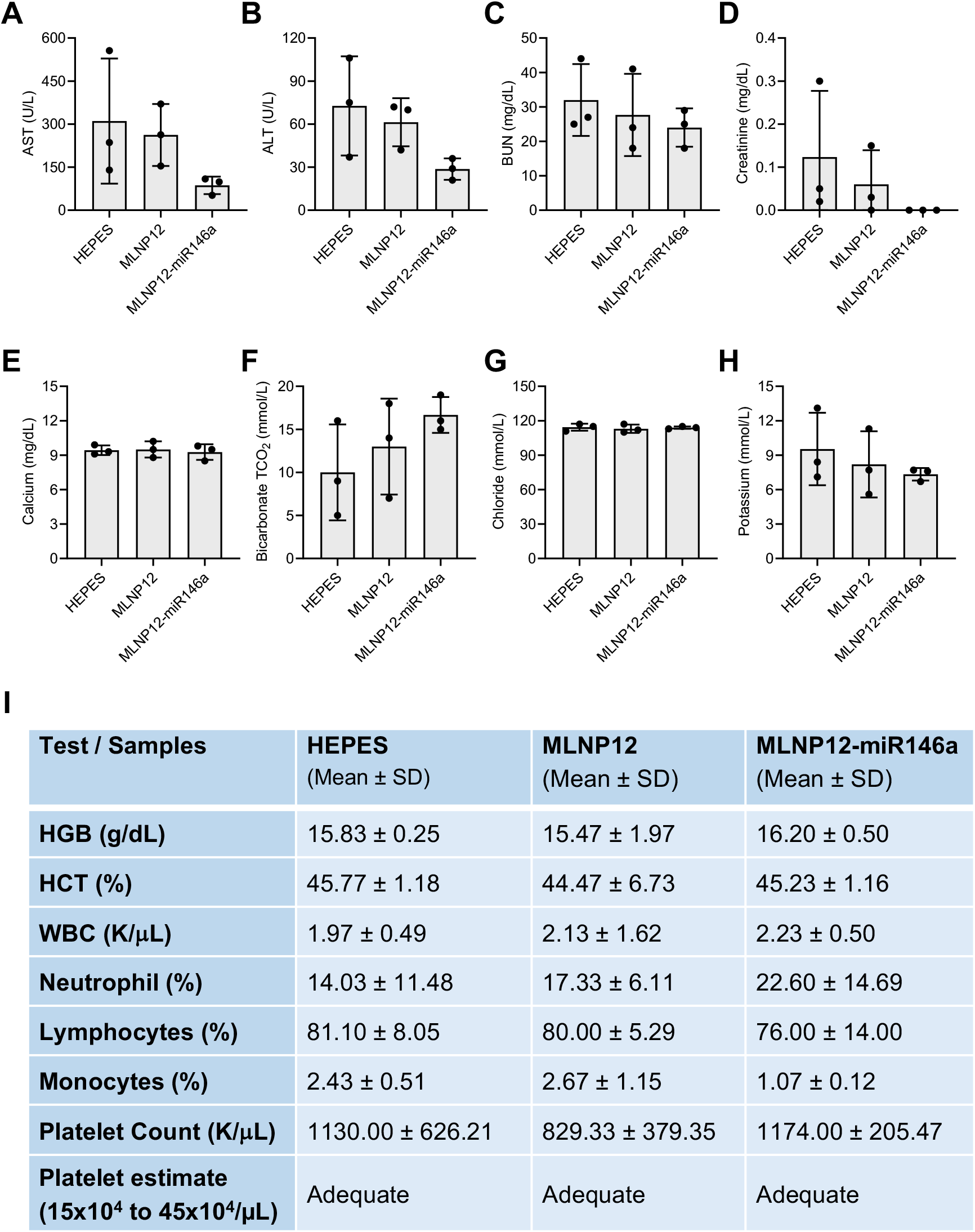
Systemic biocompatibility of empty and miR-146a-loaded MacLNP12 in vivo. In vivo cytotoxicity was assessed in mice by measuring hepatic function markers, including AST **(A)**, and ALT **(B)**, as well as renal function markers, including BUN **(C)** and creatinine **(D)**. Serum electrolytes - calcium **(E)**, bicarbonate **(F)**, chloride **(G)** and potassium **(H)** were also measured. **(I)** Hematological parameters were analyzed using whole blood samples. All experiments were performed in triplicates, and data are presented as the mean ± standard error of the mean.

Renal function was assessed by measuring blood urea nitrogen (BUN) and creatinine levels. BUN values in all treatment groups ranged from 20-40 mg/dL, consistent with normal kidney function and indicating no detectable nephrotoxicity (Figure 7C). Similarly, serum creatinine levels remained between 0.0 and 0.12 mg/dL across all groups, further confirming normal renal function (Figure 7D). Overall, AST, ALT, BUN, and creatinine levels were lower in MacLNP12-miR146a-treated mice compared with HEPES- and MacLNP12-treated mice.

Metabolic electrolyte levels, including calcium, bicarbonate, chloride, and potassium, were also evaluated. No abnormal elevations or reductions were observed, and all values remained within normal physiological ranges (Figures 7E-H).

Hematological parameters were analyzed using anticoagulated whole blood and were comparable among all groups (Figure 7I). In HEPES-treated mice, hemoglobin (HGB), hematocrit (HCT), WBC count, neutrophils, lymphocytes, monocytes, platelet count, and platelet estimate were 15.83 ± 0.25 g/dL, 45.77 ± 1.18%, 1.97 ± 0.49 K/μL, 14.03 ± 11.48%, 81.10 ± %, 2.43 ± 0.51%, 1130.00 ± 626.21 K/μL, and adequate, respectively. Corresponding values in MacLNP12-treated mice were 15.47 ± 1.97 g/dL, 44.47 ± 6.73%, 2.13 ± 1.62 K/μL, 17.33 ± 6.11%, 80.00 ± 5.29%, 2.67 ± 1.15%, 829.33 ± 379.35 K/μL, and adequate. Similarly, MacLNP12-miR146a-treated mice exhibited HGB, HCT, WBC count, neutrophils, lymphocytes, monocytes, platelet count, and platelet estimate values of 16.20 ± 0.50 g/dL, 45.23 ± 1.16%, 2.23 ± 0.50 K/μL, 22.60 ± 14.69%, 76.00 ± 14.00%, 1.07 ± 0.12%, 1174.00 ± 205.47 K/μL, and adequate platelet levels, respectively.

Collectively, these results demonstrate that both empty MacLNP12 and MacLNP12-miR146a exhibit excellent in vivo biocompatibility, with no evidence of hepatic, renal, metabolic, or hematological toxicity, supporting their suitability for therapeutic applications.

## Discussions

The advancement of microRNA-based therapeutics has been limited by poor nucleic acid stability and inefficient intracellular delivery in vivo. In this study, we engineered a four-component, macrophage-directed MacLNP system optimized for delivery of miR-146a, a key endogenous regulator of inflammatory signaling. The performance of this platform is rooted in rational lipid design and formulation strategy. The lipid-injection approach yielded particles with narrow size distribution and strong physicochemical stability across a wide pH spectrum (2.5-8) and protection against serum-associated degradation-key properties for withstanding endosomal acidification and systemic circulatory stress.

Each lipid component was selected to fulfill a defined functional role. DOTAP provided electrostatic complexation of miR-146a and protection against nuclease degradation. PA-PEG3-mannose enabled macrophage targeting via CD206 engagement while also improving colloidal stability and minimizing aggregation (11-13). Importantly, incorporation of DOPE together with linoleic acid created a membrane-destabilizing environment that likely enhanced endosomal membrane fusion and cytosolic release (11-13). This combinatorial lipid architecture appears to underlie the efficient intracellular delivery in primary macrophages and functional target engagement observed in our studies.

MicroRNAs function as endogenous network regulators controlling macrophage activation, inflammatory amplification, and fibrotic remodeling (14-38). Among these, miR-146a acts as a critical negative-feedback regulator of innate immune signaling (25-29). Our previous work established a clear inverse relationship between miR-146a levels and the severity of FBR: increased miR-146a suppresses macrophage accumulation, foreign body giant cell formation, and collagen deposition, whereas genetic deletion exacerbates these pathological features (20). Thus, therapeutic restoration of miR-146a represents a mechanistically grounded strategy to attenuate biomaterial-associated inflammation.

In the present study, MacLNP-miR146a demonstrated prolonged in vivo persistence, restricted off-target biodistribution, and sustained retention at the administration site. Both localized and systemic delivery studies confirmed targeted accumulation with minimal unintended organ distribution beyond expected hepatic clearance. Furthermore, comprehensive biochemical and hematological analyses revealed no evidence of hepatic, renal, metabolic, or hematologic toxicity, supporting favorable in vivo tolerability of both empty and miR-loaded formulations. Despite these encouraging findings, certain limitations warrant consideration. While short-term biodistribution and safety profiles were favorable, extended longitudinal studies will be necessary to fully define long-term clearance kinetics, potential delayed accumulation, and chronic safety. Additional evaluation in large-animal models and implant-relevant disease contexts will further strengthen translational positioning.

Overall, our findings support MacLNP-miR146a as a stable, targeted, and biologically effective delivery platform for controlling inflammation associated with biomaterial implantation. By integrating rational lipid design, macrophage specificity, efficient endosomal escape, and scalable manufacturing, this system addresses key barriers that have historically constrained microRNA therapeutics and provides a promising foundation for future clinical translation.

## Acknowledgements

This work was supported by an NIH (R01EB024556) grant to Shaik O. Rahaman. We acknowledge the DLAR Imaging Core Facility, located at A.J. Clark Hall at the University of Maryland, College Park for providing imaging equipment assistance and consultation.

## Authors contributions

SOR and MIK conceived the study, designed the experiments, analyzed the data, and wrote the manuscript. MIK and KRS performed the experiments. SOR edited the manuscript. All authors reviewed and approved the final content of the manuscript.

## Data availability

All data generated and used during this study are included in this article.

